# Micro-Cm: restrictase-free microbiome-wide chromosome conformation profiling

**DOI:** 10.64898/2026.09.11.750865

**Authors:** Marcos Bermejo-Ruiz, Carolin Wilhelm, Heike Budde, Ruth E. Ley, Alexander V. Tyakht

## Abstract

Mobile genetic elements like plasmids, viruses and transposons can considerably augment the genomic repertoire of individual bacterial members of a complex multi-species microbiome and influence community dynamics. As linking a mobile element to its bacterial host based on metagenome sequencing alone proves challenging, such assays have been augmented with high-throughput chromosome conformation capture (Hi-C). However, the efficacy of Hi-C metagenomics is constrained by the protocol limitations and a lack of ground-truth reference datasets. In order to overcome these limitations, we present Micro-C metagenomics (Micro-Cm) - an adaptation of a superior, restrictase-free Micro-C technique for processing microbiome samples and mapping plasmid-host associations. We validated the developed experimental protocol and bioinformatic workflow on a simulated, defined consortium of diverse gut bacterial species and applied them to a long-read human gut microbiome sample. The proportion of valid reads in the synthetic community was an order of magnitude higher than that observed in multiple Hi-C metagenomic studies. For both samples, we obtained high-quality contact maps, which in the case of the synthetic community revealed fine-scale chromosome interactions. Moreover, successful recovery of plasmid-host interactions in the simulated community validated the method, which we then applied to the real stool sample. Our plasmid-host association analysis in a complex bacterial community successfully identified bacterial hosts for most of the identified complete plasmids. Our results show that Micro-Cm method improves profiling of complex microbiomes, exploration of mobile genetic element dynamics and community-wide, detailed investigation of chromosomal conformation patterns.

## Introduction

Metagenomics is a widely used approach for characterizing the taxonomic composition and functional potential of complex microbial communities. Part of this functional diversity is associated with plasmids, which are extrachromosomal genetic elements that carry a broad range of genes. In addition to genes required for their replication and maintenance, plasmids frequently encode accessory functions that influence bacterial adaptation and ecological interactions, including antibiotic resistance genes (McInnes et al. 2020) or defense systems (Stockdale and Hill 2023). To understand the transmission and dynamics of these microbiome-wide accessible genetic pools, it is crucial to link the plasmids to their bacterial hosts - a task that remains a substantial methodological challenge (Yang et al. 2025).

Building upon metagenomic plasmidome profiling, multiple techniques have attempted to link plasmids to their hosts. Sequence similarity methods struggle due to the plasticity of plasmid sequences. Hybrid lab-computational techniques, such as the high-fidelity targeted ProFiT-SPEci-FISH (Zorea et al. 2025), are effective for determining the host of a specific plasmid, but cannot be applied for broad plasmid-host linking. Hi-C, a chromatin conformation capture technique, has been integrated into metagenomic workflows. It estimates the three-dimensional spatial proximity of DNA fragments, independent of their linear sequence context. By doing so, Hi-C not only refines contig binning and facilitates the recovery of higher-quality metagenome-assembled genomes (MAGs) (Burton et al. 2014), but also enhances the capacity to link mobile genetic elements to their host organisms (Stalder et al. 2019). These improvements are due to the fact that proximal DNA is expected to exhibit a higher Hi-C signal, whereas DNA belonging to different bacterial species (and thus not co-occurring in the same cell) is anticipated to show significantly lower or no signal. When applied in clinical settings, this approach has shown potential in deeply evaluating drug resistance genetic determinants and other risk factors associated with opportunistic microbes (Ivanova et al. 2021; Revel-Muroz et al. 2026).

While Hi-C metagenomic data offer a layer of biological information that is largely inaccessible to conventional metagenomics, the original site-specific restrictase-based Hi-C metagenomic protocol poses considerable limitations. First, its resolution is limited: genomic regions lacking restriction sites are not cleaved, making it difficult to obtain high-resolution chromosomal interaction maps. Second, in the metagenomic context, only a small fraction of sequencing reads represent valid ligation products that convey the interaction signal, a phenomenon that has been reported consistently across multiple studies (Sonets et al. 2024; Regmi et al. 2025; Rojas et al. 2023; Wu et al. 2023). Third, the absence of reference data from well-characterized MGE-augmented microbial consortia has impeded comprehensive benchmarking of the different steps required for Hi-C metagenomics, including filtering and normalization procedures (Shatadru et al. 2025). Additionally, constructed synthetic communities exhibit limited phylogenetic diversity among bacterial species and their associated plasmids (Beitel et al. 2014).

Improved versions of the original Hi-C method have been introduced trying to overcome the previously mentioned limitations and have been successfully applied primarily for eukaryotic organisms, including DNAse Hi-C (Ma et al. 2015), Hi-C 3.0 (Lafontaine et al. 2021), Pore-C (Deshpande et al. 2022) and Micro-C (Hsieh et al. 2015). Micro-C has been applied to bacterial isolates to obtain unprecedentedly detailed chromosome conformation maps and to discover 3D genomic patterns in *Escherichia coli* (Gavrilov et al. 2025). It differs from Hi-C primarily in one aspect - the use of micrococcal nuclease instead of a restriction enzyme. Micrococcal nuclease cuts DNA randomly at virtually any position where the DNA is accessible. As a result, DNA fragments have more uniform sizes and the resulting resolution is increased. Despite offering clear advantages over Hi-C, this method has not yet been applied to a complex microbial community. Here, we further adapt this technique for complex bacterial communities. We tested the method with an MGE-augmented simulated community composed of five different plasmid-containing species and demonstrated the remarkable performance of the protocol on a real human stool sample.

## Materials and methods

### Simulated community

#### Bacterial cultivation

We selected a phylogenetically diverse set of five bacterial species representative of the human microbiome (*Bacteroides fragilis, Collinsella aerofaciens, Enterococcus faecium, Prevotella nigrescens* and *Shigella flexneri*), using one strain of each to construct a synthetic (“mock”) defined community (Supplementary Table 1). The initially estimated number of replicons per species varied, with individual species containing up to two chromosomes and four plasmids. Each bacterial species was cultured overnight anaerobically (95% N_2_ and 5% CO_2_ atmosphere) at 37 °C in modified PVG medium (see Suppl. Methods). After 48 h of incubation, the optical density at 600 nm (OD_600_) was measured (Supplementary Table 1). Two independent replicates of the mock community were prepared. For each replicate, 400 µL of each culture was combined and centrifuged at 4,000 × g and 4 °C for 5 min. The supernatant was discarded, and another 400 µL of each species culture was added to the pellet. The combined cell suspension was resuspended in 1 mL of fixing solution (3 % formaldehyde in 0.9 % NaCl) and incubated at room temperature for 30 min, followed by 30 min on ice. Subsequently, 66.7 µL of 2 M glycine was added and the cells were centrifuged at 17,000 × g and 4 °C for 10 min. The pellet was re-suspended in 100 µL PBS. The samples were then snap-frozen in liquid nitrogen and stored at -80 °C until proceeding with the Micro-Cm protocol.

#### Long-read sequencing of bacterial isolates

To ensure that the genomes from the laboratory-derived strains did not diverge substantially from their corresponding reference genomes deposited in the NCBI Refseq database, each strain was sequenced prior to pooling. Long-read sequencing (with each bacterial isolate sequenced separately and assigned a unique barcode) and quality filtering of reads were performed as described below. Genome assemblies were generated using Autocycler v0.5.2 (Wick et al. 2025). Although this tool failed to recover the short plasmids of *P. nigrescens*, they were successfully identified by directly aligning the reads to the reference sequences. Additionally, two newly acquired plasmids were detected in *S. flexneri* and were thus included in the analysis.

To re-arrange the assembled sequences so that their genomic start positions matched those of the reference genomes, the tools SeqKit v2.8.0 (Shen et al. 2024) and SAMtools v1.22.1 (Danecek et al. 2021) were used. The re-arranged assemblies were compared against the reference sequences using MUMmer v3.23 (Kurtz et al. 2004). As no substantial differences were observed, modification of the reference genomes was limited to adding the newly discovered plasmids to the *S. flexneri* genome.

#### Construction of chromosomal contact maps

HiCExplorer v3.7.6 (Wolff et al. 2022) was used to generate contact matrices. First, the Micro-Cm paired-end reads (R1 and R2) were aligned independently to the reference genome using BWA v0.7.19-r1273 (Li and Durbin 2009). Then a contact matrix was constructed using the `hicBuildMatrixMicroC` command (tested across a range of resolutions to define the optimal one) that also outputs the number of valid Micro-Cm contacts. A histogram of contact-intensity distribution was produced with the `hicCorrectMatrix diagnostic_plot` command, and the threshold automatically suggested by the tool was applied to normalize the matrix using `hicCorrectMatrix correct`. Finally, the normalized matrix was visualized with the `hicPlotMatrix` command. Each heatmap cell reflects the estimated interaction intensity between the respective DNA fragments of a specified fixed length (resolution).

#### Simulation of a fragmented metagenome assembly from reference genomes

To assess the performance of the proposed method in associating plasmids with their bacterial hosts, we simulated a fragmented metagenomic assembly by randomly splitting the reference genomes into “contigs”. Multiple iterations (n = 25) were performed, in which all the chromosomes and the sequences of two-thirds of the plasmids were split with varying fragmentation levels. The remaining one third of the plasmid sequences were kept intact (to simulate a real scenario in which some of the plasmids could be fragmented but some others will be retrieved as full circular sequences). NormCC (Du and Sun 2023) was used to calculate the normalized contact matrix, and the chromosome-plasmid interactions were evaluated as described below. The results were averaged across the iterations.

#### Summarizing the spatial proximity of a replicon along another replicon

To characterize the spatial distribution of a selected replicon (query) along another replicon (target) - e.g., for a chromosome–chromosome or plasmid–chromosome pair - both sequences were divided into bins, and for each target bin, its contact intensities with each query bin were summed. This value was then normalized by dividing it by the total number of query bins. For plasmid–chromosome comparisons, an additional normalization step was applied. Specifically, the 98th percentile and 2nd percentile of the contact distribution were used as the upper and lower reference values, respectively to remove contact outliers that could bias the normalization. Contact frequencies were then rescaled to a range between 0 and 1 based on these percentile thresholds.

For the comparison of the two-chromosome species (*P. nigrescens*), the origin of replication was determined as follows: for chromosome 2, the origin was identified based on the GenBank annotation; for chromosome 1, the putative origin region was predicted using the Ori-Finder 2022 web service (Dong et al. 2022), since no origin of replication was annotated. Gene annotations from RefSeq were used, and gene functional classes were assigned according to the BV-BRC database (Olson et al. 2023).

### Human gut microbiome

#### Sample collection

A stool sample was collected from an anonymous human donor. The sample collection was approved by the local ethics committee (Ethik-Kommission an der Medizinischen Fakultät der Eberhard-Karls-Universität und am Universitätsklinikum Tübingen, protocol number: 456/2023A). The sample (ID: MC1) was kept on ice and frozen at -80 °C three hours after the collection and then processed the next day.

#### Short- and longread metagenome sequencing

Genomic DNA extraction and library preparation for Illumina short-read sequencing were performed according to the protocol described in (Suzuki et al. 2022).

For Nanopore long-read sequencing, DNA was extracted from the stool sample using the Qiagen PowerFecal Pro Kit. Cell lysis was achieved by vortexing the sample at maximum speed for 10 min, and then the elution was performed in 50 µL of nuclease-free water. Beadbased cleanup was performed using 0.45× AMPure XP beads: the beads were added to the eluate, and the mixture was incubated for 5 min at room temperature before being placed on a magnetic rack for 5 min until the solution cleared. The pellet was washed twice with 80% ethanol, dried for 2 min, and the DNA was eluted in 40 µL of nuclease-free water. Sequencing libraries were prepared using the Oxford Nanopore Technologies Native Barcoding Kit 24 V14. Three distinct barcodes were used for the same sample, with each receiving 1000 ng of input DNA. The final concentration of the library was 13 ng/µL, corresponding to a total of 300 ng (50 fmol).

#### Long-read metagenome assembly

A scheme summarizing the steps required for the assembly of the stool sample reads is shown in Supplementary Fig. 1. In brief, Chopper v0.9.0 (De Coster and Rademakers 2023) was used to remove low-quality Nanopore reads (quality score < 9 or length < 900 bp). The filtered reads were assembled using Flye v2.9.5 with the `--meta` flag (Kolmogorov et al. 2020). The resulting contigs were binned into MAGs using three conventional (Micro-Cm-independent) tools: MetaBAT2 v2.17 (Kang et al. 2019), SemiBin2 v2.2.1 (Pan et al. 2023) and MaxBin2 v2.2.7 (Wu et al. 2016).

In parallel, the contigs were also binned using tools that leverage the Micro-Cm data. For this purpose, the Micro-Cm reads were aligned to the contigs using BWA-MEM (Li and Durbin 2009) with the `-5SP` argument. A normalized matrix of contacts was obtained using a custom version of NormCC (part of MetaCC) (Du and Sun 2023), which was modified to enable restrictase-free data processing. This matrix was used as an input for two binning tools - ImputeCC v1.1.0 (Du et al. 2024) and MetaCC v1.2.0 (Du and Sun 2023) - to generate one set of MAGs (a binning) per tool.

These two Micro-Cm-based binnings, along with the three Micro-Cm-independent binnings, were aggregated and dereplicated using DAS Tool v1.1.7 (Sieber et al. 2018). Taxonomic classification of the final MAG set was performed using GTDB-Tk v2.1.1 based on GTDB taxonomy release 207 (Chaumeil et al. 2022), and MAG quality was assessed with CheckM2 v1.0.1 (Chklovski et al. 2023)).

#### Identification of circular plasmid contigs

Illumina short reads were assembled using metaSPAdes v4.2.0 (Nurk et al. 2017). The resulting assembly graphs were processed with metaplasmidSPAdes v4.2.0 (Antipov et al. 2019) and SCAPP v0.1.1 (Pellow et al. 2021) to identify small circular sequences potentially representing plasmids. Each of these sequences was classified using ViralVerify v1.1 (Antipov et al. 2020) into one of three categories: chromosomal, viral, or plasmid; only the plasmid sequences were retained. The metaplasmidSPAdes- and SCAPP-based plasmid sequence sets were dereplicated using MMseqs2 v18.8cc5 (Steinegger and Söding 2017) to yield a nonredundant collection of plasmids. For our proof-of-principle analysis, and to focus on the highquality data, we did not consider fragmented, linear plasmid-like contigs.

#### Construction of chromosomal contact maps

The contact map was visualized using R. In this case, we constructed a contig-level map in which each of the pixels represent each of the contigs belonging to a MAG. Only the contacts whose intensities were above the threshold and only the high-quality MAGs were shown.

### Micro-Cm sequencing

The Micro-Cm protocol described below was adapted from the Micro-C protocol from (Gavrilov et al. 2025), with full details provided in the Supplementary Methods. Briefly, DNA was crosslinked in two steps: first with formaldehyde, followed by disuccinimidyl glutarate (DSG) - a step not present in the Hi-C protocol. Following the fixation, cells were disrupted to free the DNA, and the DNA was digested using micrococcal nuclease (MNase) rather than the restriction enzyme(s) used in Hi-C. After MNase digestion, dephosphorylation was done (another step absent in Hi-C), and the fragmented ends were filled with biotinylated nucleotides, and the DNA fragments were ligated. The biotin from the unligated ends was removed, and crosslinks were reversed (steps performed post-DNA extraction in standard Hi-C). The DNA was extracted using bead purification, and RNA was digested. The purified DNA was then sonicated to shear it into shorter fragments. To isolate the ligated fragments of interest, a biotin pull-down was performed, followed by A-tailing for sequencing preparation. Finally, sequencing adapters were ligated, PCR amplification was performed, and the resulting libraries were sequenced on an Illumina platform. For the simulated community, the two replicates were sequenced separately, and the resulting datasets were combined for further analysis.

### Secondary analysis of Micro-Cm contacts

For Micro-Cm-generated contacts, the noise-to-signal ratio was calculated based on chromosomal read counts between and within genomes or MAGs (depending on the sample type), as described in (Revel-Muroz et al. 2026). The proportion of valid Micro-Cm reads was determined using quality measures from HiCExplorer. Specifically, the number of valid Micro-Cm reads was divided by the total number of Micro-Cm reads. To account for the incomplete representation of the metagenome by the genomes, this value was multiplied by the ratio of the total reads to mappable reads. For the real stool sample, this ratio was calculated after filtering out the short metagenomic contigs. (To evaluate the impact of short contigs on the final proportion of valid Micro-Cm reads, this procedure was repeated across various minimum contig length thresholds.) After the normalization, the contact matrix represented the normalized contact intensities for each pair of contigs in the metagenome. Based on the previously described allocation of contigs into the final sets of Micro-Cm-independent and - dependent MAGs, each contact was classified as either “between-MAG” or “within-MAG”. We compared the distribution of the generally weaker “between-MAG” contact intensities with the stronger “within-MAG” intensities to define an optimal threshold for distinguishing true signal (within-cell contacts) from noise (spurious/between-cell contacts). A precision-recall (PR) analysis was conducted on these intensities using the PRROC R package, allowing us to statistically determine the optimal threshold as the value maximizing the F1 score. All contacts falling below this threshold were discarded. Contacts linking two plasmid contigs were recorded directly to represent spatial associations between plasmids. To quantify the interaction intensity between a plasmid contig and a MAG, we calculated the median contact intensity between the plasmid contig and all contigs comprising that MAG.

## Results

### Simulated community

Processing the high-coverage sequencing data from the mock community (350 million reads) revealed that 31.2% of them were valid Micro-Cm reads (i.e., properly aligned and containing a ligation event). This proportion was considerably higher than the average of those reported in several previous Hi-C-based microbiome assays (Stalder et al. 2019; Beitel et al. 2014; Rojas et al. 2023; Sonets et al. 2024; Revel-Muroz et al. 2026) and comparable to the yield obtained in the pure *E. coli* Micro-C experiment (Gavrilov et al. 2025). The contact noise-to-signal ratio (see Methods) was very low (0.04) (Revel-Muroz et al. 2026), further confirming the high quality of the generated data. Together with availability of the complete genomes, this provided a solid foundation for a multi-level investigation of 3D genome organization within each member species.

The pan-community chromosomal contact map (1 Kbp resolution; Fig. 1) featured five prominent diagonal squares corresponding to each bacterial species, clearly illustrating the prevalence of the within-species over between-species contacts. Leveraging the enhanced resolution provided by the Micro-Cm approach, we explored the limits of contact map construction. For each species, we successfully generated highly complete maps at a resolution as fine as 20 bp. These maps revealed fine-scale chromatin patterns (Supplementary Fig. 2), particularly resembling those discovered in the *E. coli* Micro-C study (Gavrilov et al. 2025), warranting future in-depth investigation.

**Fig 1.**
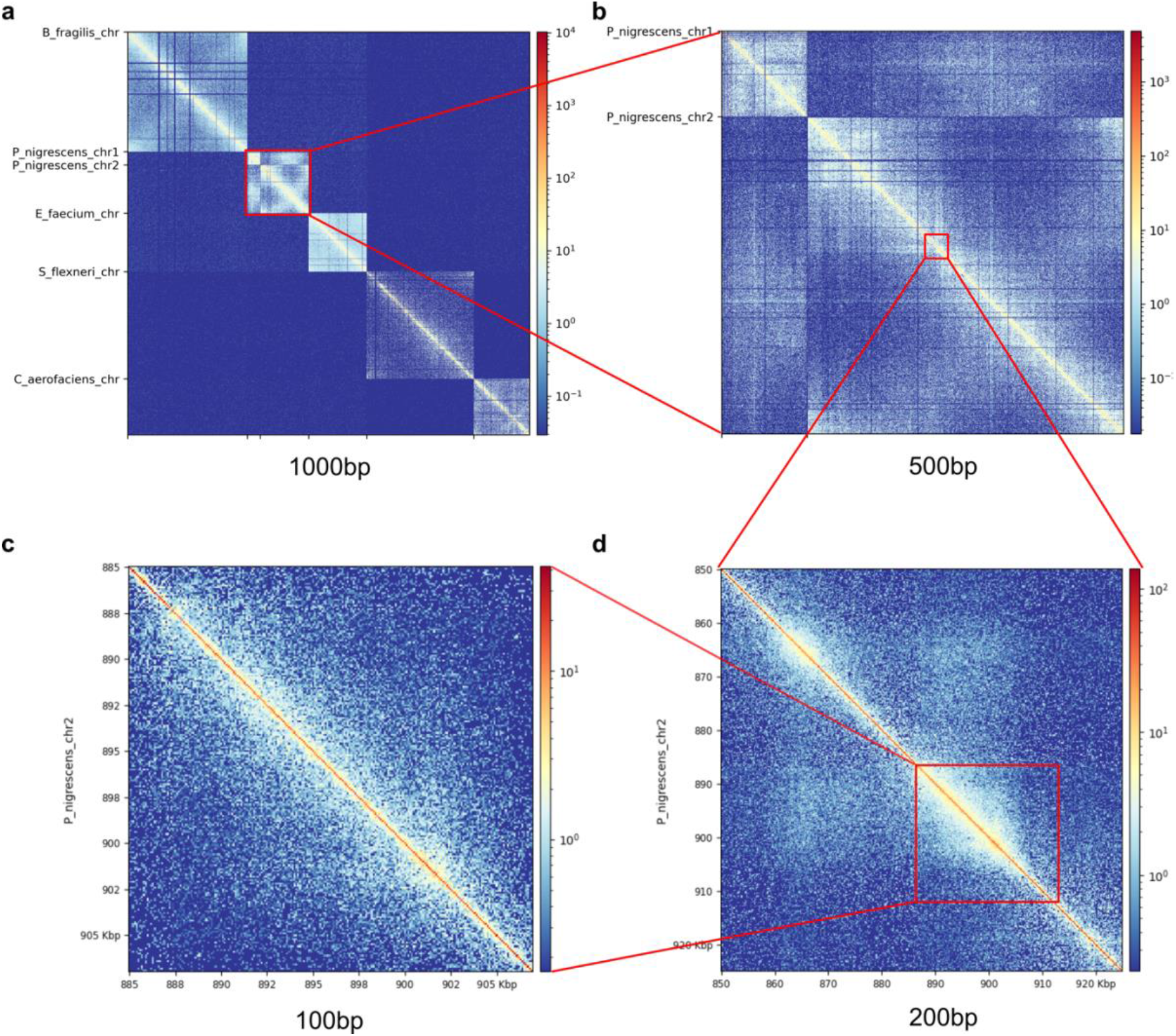
Contact maps at increasing resolutions observed in the simulated community. The resolution of the map is specified at the bottom of each panel. The increasing resolution allows visualization of the entire community (a), interactions between the two chromosomes of one community member (*Prevotella nigrescens*) (b), and different regions within one bacterial chromosome (*P. nigrescens* chr2) (c and d).

At a more granular level, we investigated the contacts of individual replicons within each species. We observed a notable lack of contacts involving the four *P. nigrescens* plasmids. Further inspection of the standard genomic (non-Micro-Cm) sequencing data revealed that each of these plasmids had an order of magnitude lower read coverage than the *P. nigrescens* chromosomes (Supplementary Table 2). This finding aligned with the observation that their sequences mapped perfectly to various chromosomal regions, strongly suggesting plasmid integration. Micro-Cm contacts were also undetected for the single *B. fragilis* plasmid; however, the underlying cause was different. A comparison of the sequencing datasets suggested that this plasmid was likely lost during the cultivation of the isolate prior to pooling it with the other species cultures to form the mock community. Consequently, these plasmids were excluded from further analysis, leaving seven plasmids in the further analysis.

The presence of species with multiple replicons allowed us to explore their intra-cellular co-localization. For *P. nigrescens*, we discovered that its two chromosomes were spatially distributed in an asymmetric manner. Specifically, the smaller chromosome 1 preferentially associated with the 0.8-1.7 Mbp region of chromosome 2 (Fig 2a), with the distribution peak roughly coinciding with the replication origin of chromosome 2. Interestingly, the distribution of chromosome 2 contacts along chromosome 1 was much more uniform (Fig 2b). A functional overview of the region on chromosome 2 that was highly proximate to chromosome 1 revealed a particular increase of the ribosome biogenesis genes ((Supplementary Table 3), likely linked to the presence of the replication origin in that region (Soler-Bistué et al. 2015).

**Fig 2.**
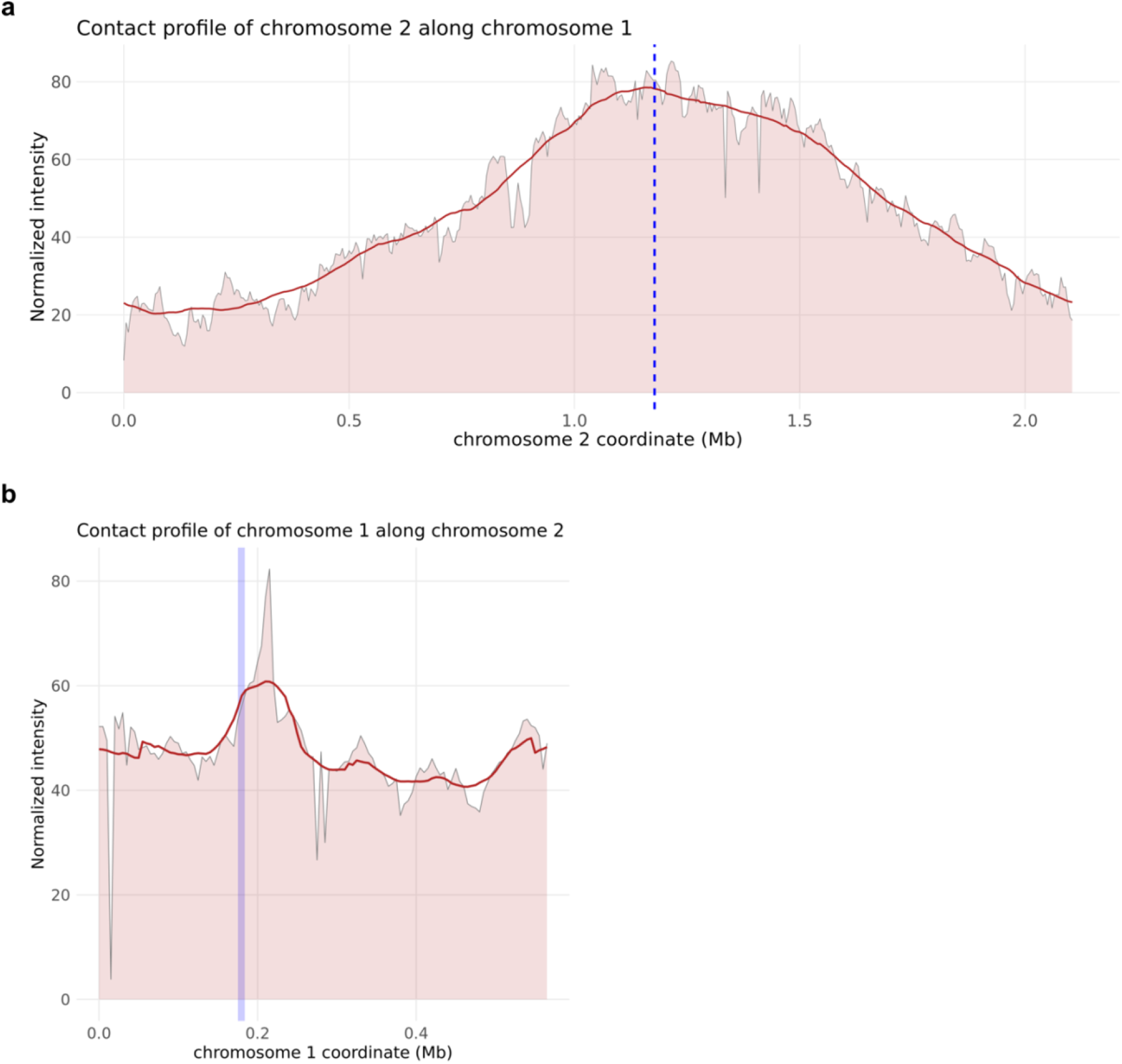
Summarized spatial co-localization of the two chromosomes of Prevotella nigrescens recovered from the simulated bacterial community. Resolution: 5 Kbp. Vertical blue lines denote replication origins based on the range of predicted origins (chr1) and GenBank annotation (chr2). a) Contact profile of chr2 along chr1. b) Contact profile of chr1 along chr2.

We summarized the distribution of contact intensities as a function of genomic distance by plotting contact scaling curves (Fig 3a). All chromosomes exhibited a near-inverse log-linear dependence - a hallmark of DNA organization universally observed across domains of life (Halverson et al. 2014) - and the contacts for plasmids larger than 25 Kbp scaled similarly to their respective chromosome. Because the analyzed bacterial replicons were circular, a characteristic uptick in contact intensity was observed at the tail end of every curve. Excluding the three >5 Kbp plasmids, we calculated the slopes of the log-scaled contact curves for each chromosome and <5 Kbp plasmid. These slopes, which visually clustered into 2 groups, ranged from -1.0 and -0.5. This is generally consistent with the value previously reported for the *E. coli* chromosome (Wasim et al. 2021) and reflects that the spatial complexity of bacterial genomes does not perfectly conform to either equilibrium or a fractal globule models. Conversely, the three small plasmids exhibited less steep slopes, presumably reflecting a higher degree of global compaction across the entire replicon.

**Fig 3.**
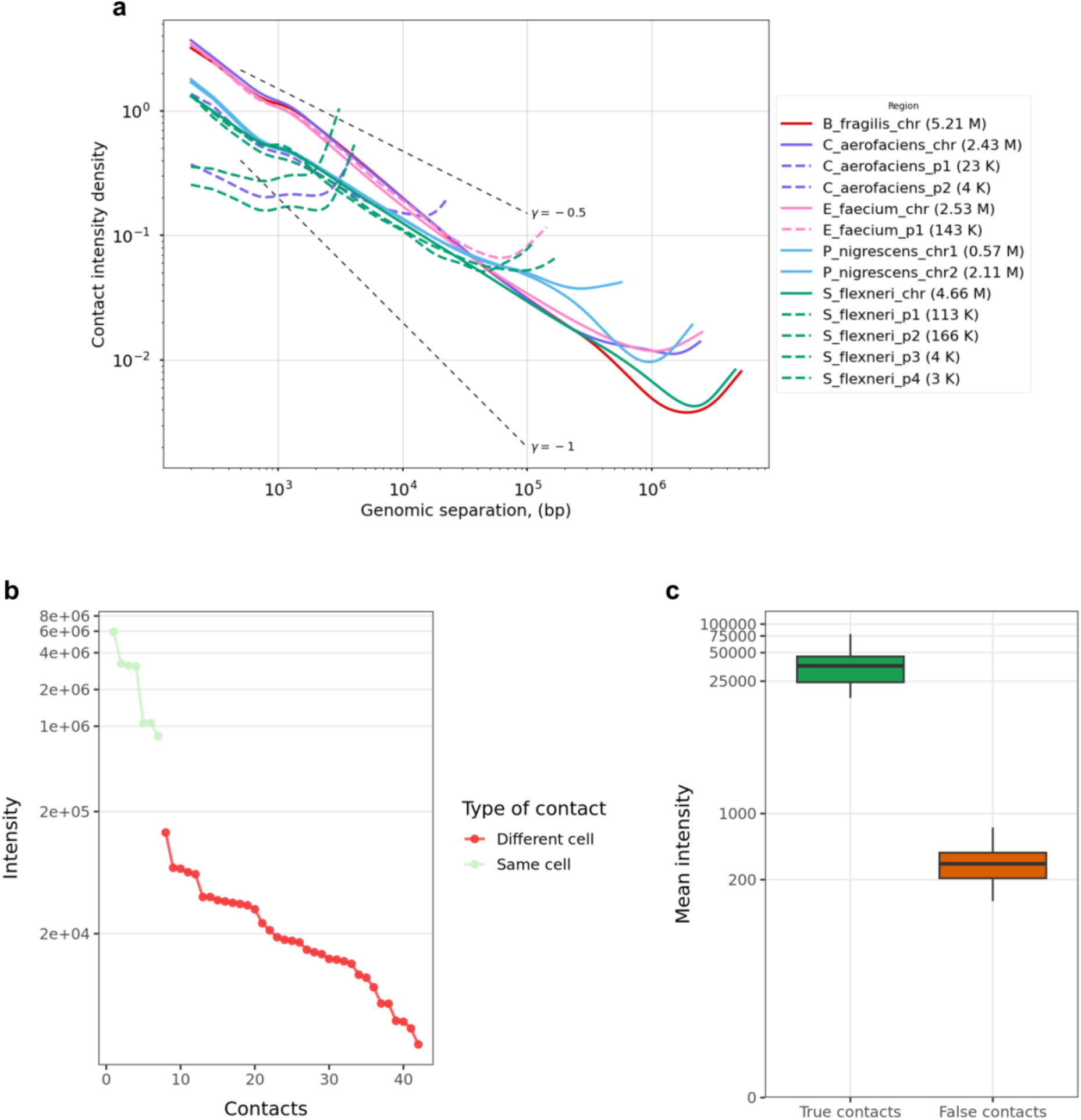
Statistics of the characteristics of plasmid contacts in the simulated community. a) Scaling curves representing contact densities as a function of genomic distance. Sequences belonging to the same species share the same color. Solid lines represent chromosomes, and dashed lines represent plasmids. An increase in interaction intensity is observed at the end of each sequence (chromosomes and plasmids), reflecting sequence circularity. b) Representation of the contact intensities between the replicons in descending order. Contacts between a plasmid and its host chromosome are shown in green, while contact between a plasmid and other chromosomes are shown in red c) Comparison of the median values of true and false contacts across 25 iterations of simulated metagenomic contigs derived from the reference genomes.

When comparing inter-replicon contact intensities across the bacterial species, the top 16.7% most intense plasmid-chromosome pairs captured all of the biological ground-truth associations (2.6 ± 1.8 × 10^6^, mean ± s.d.). In contrast, all non-existent (false) associations exhibited substantially lower intensities (27.0 ± 27.0 × 10^3^). On average, true contacts showed 97-fold higher intensities than the false contacts (Fig 3b), demonstrating that Micro-Cm is a highly accurate approach for linking plasmids to their bacterial hosts. Similar results were obtained when the reference genomes were randomly fragmented *in silico*, indicating that the approach remains robust with the imperfect assemblies typical of real-world metagenomic studies (Fig 3c).

### Human gut microbiome

For the human stool sample, deep coverage data were generated using both long-read WGS (17 million reads) and short-read Micro-Cm sequencing (356 million reads). The proportion of valid Micro-Cm reads was 10.3%, which is higher than reported in earlier Hi-C studies of this sample type; the estimate remained stable even when short contigs were removed across various sequence length thresholds (Supplementary Fig. 3). The generated data allowed us to reconstruct a large number of MAGs (n = 175), of which 162 received a species-level taxonomic assignment and 101 had high quality (>90% completeness, <5% contamination; Supplementary Table 4). Among the latter, 64 were derived using Micro-Cm data, and 37 via conventional binning. As with the simulated community described above, the Micro-Cm noise-to-signal ratio was very low (0.024).

The quality and distribution of the Micro-Cm contacts were initially evaluated visually via the contig-level contact map (Supplementary Fig. 4). As expected for Micro-Cm data, contigs belonging to the same MAG had more intense contacts than those belonging to different MAGs (intra MAG contacts: 49% values are zero, median of the non-zero values: 16.4 × 10^3^, 95% CI: 16.0 - 17.0 × 10^3^, inter-MAG contacts: 99.8% zeros, median of non-zero values: 4.9 × 10^3^, 95% CI: 4.8 - 5.0 × 10^3^; Wilcoxon rank-sum test, p < 2.2 × 10^−16^). The contact distribution also followed the expected pattern (Fig 4a). Between-MAG contacts showed weaker intensities compared to within-MAG contacts, allowing for a clear distinction between the distributions using the calculated threshold (see Methods).

**Fig 4.**
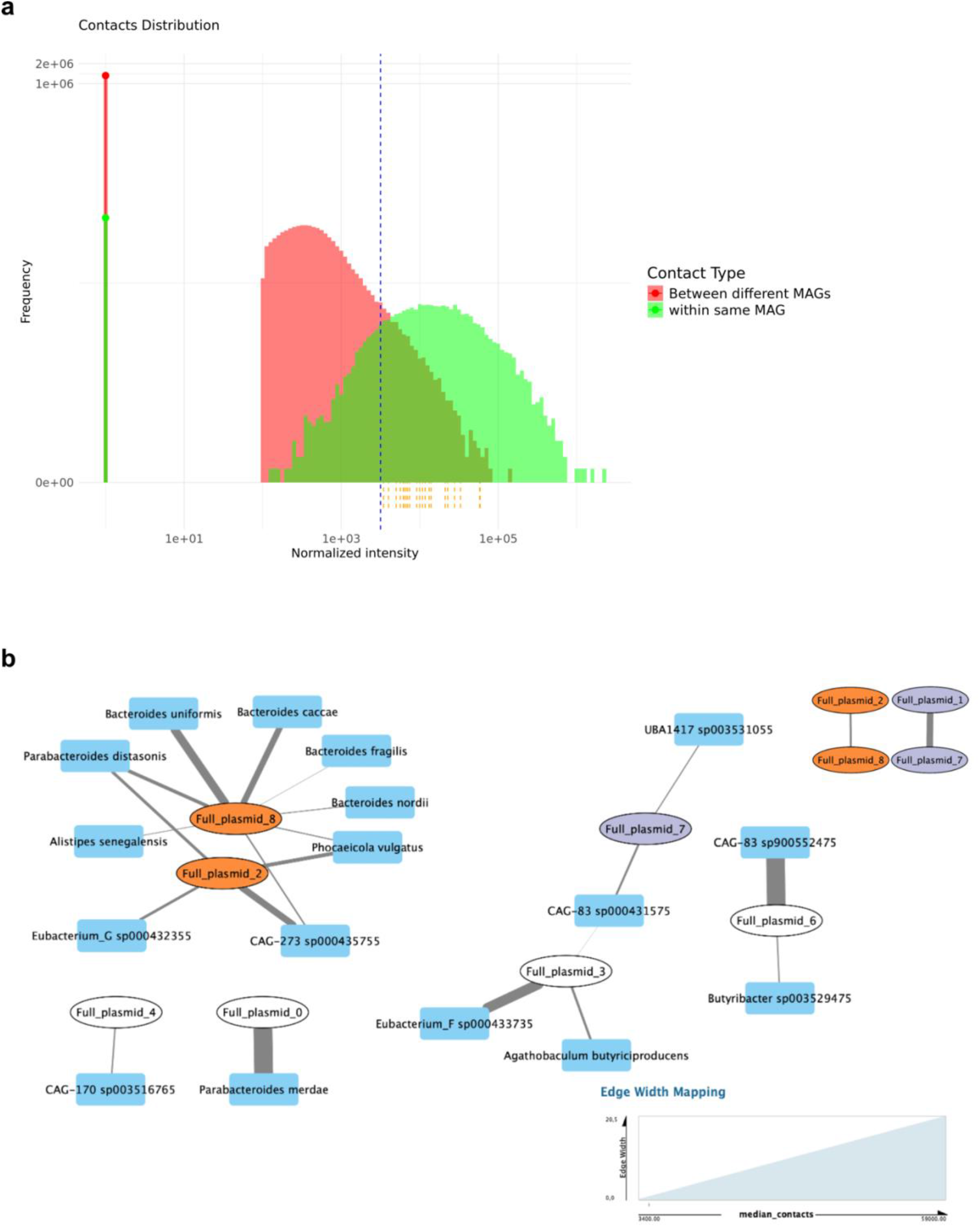
Micro-Cm of a human gut microbiome sample. a) Distribution of normalized contact intensities within and between MAGs. b) Network of contacts between plasmid contigs and other plasmid contigs (top right), as well as between plasmid contigs and MAGs (the rest). MAGs are represented as blue rectangles, while plasmid contigs are shown as ellipses. The color of the plasmid contigs indicates their contacts with other plasmid contigs; plasmid contigs lacking inter-plasmid contacts remain white. MAG-MAG contacts are not shown.

Nine complete plasmids (labeled “0”-”8”) were assembled from the stool metagenome. Following the methodology established for the simulated community, we calculated the contacts between the complete plasmid contigs and the MAGs (Fig 4b). The largest hub of contacts was formed by one plasmid shared across several *Bacteroides* species, although most of the plasmids only had contacts with 1-2 bacterial hosts. In most two-host cases, one host showed a distinctly stronger interaction intensity. Alignment of the plasmid sequences against the BLAST core nucleotide database showed that most assembled sequences had at least a partial alignment with known plasmid sequences, supporting their plasmidic identity (Supplementary Table 5). Plasmid 5, for which we found no host association, did not produce any significant sequence matches in the alignment. Plasmids 0, 4 and 7 aligned mainly to chromosomes, while plasmids 2 and 8 aligned with *Bacteroides* plasmids, as expected. Finally, for plasmids 1, 3 and 6, BLAST results showed that a host bacterium had not been previously found, suggesting novel plasmids and host associations were identified.

## Discussion

We demonstrated the adaptation of the Micro-C technique to microbial communities of varying complexity. The high yield of valid data, improved resolution, and simpler experimental procedure suggest that Micro-Cm will serve as a robust alternative to Hi-C metagenomics. The experimental and data analysis components of the technique are, in principle, universal and can be applied for restrictase-independent 3D genomic analysis of microbial communities from diverse ecological niches.

We demonstrated how Micro-Cm can effectively link plasmids with their bacterial hosts. Similarly, it can be used for linking other mobile genetic elements, such as phages and transposons. The method can uncover associations in microbiomes that were previously obscured by noise or undetectable due to the bias caused by sparsely distributed restriction sites. The generated dataset is the first to provide a ground-truth Micro-Cm reference to be used for future optimization of algorithms and normalization strategies of spatial DNA linking. As a limitation, some of the initially planned plasmids had to be removed from the analysis because they had their sequences integrated into the host genomes or were discarded by the bacteria during sample preparation. Our observations indicate that even in a simple simulated community, plasmid carriage is highly volatile - plasmids can be easily lost, or, conversely, acquired. These caveats should be considered in the future studies aiming to generate 3D genomics reference sets based on more complex MGE-carrying synthetic communities. Improved strain selection and refined bioinformatic handling of sequences shared between plasmids and chromosomes can further improve the accuracy of plasmid-bacterium linking. Candidate links detected via untargeted Micro-Cm can subsequently be used to optimize the application of targeted, high-fidelity methods by narrowing the search parameter space.

Another valuable application of Micro-Cm is improving metagenomic assembly. Our results suggest that high-quality assemblies derived from long-read sequencing data provide greater advantages for use with Micro-Cm than short-read sequencing. In eukaryote-rich microbiomes, this combination can also improve the recovery of 3D genomes from eukaryotic members (e.g., yeast). The high resolution achieved will enable comprehensive investigation of not just the interactions between multiple replicons within a bacterial cell, but also the hierarchical structure of chromosomal patterns, such as small loops and chromosomal interaction domains. This is achieved by directly reconstructing the 3D structures of abundant microbiome members - not just their chromosomes, but even their plasmids and phages - without the need for cultivation or isolation. A limitation to achieving this goal is sequencing depth. The detection of fine-scale interaction patterns in our simulated community required >300 million paired-end reads; naturally occurring complex communities will likely demand higher coverage to achieve the same level of detail. Following our proof-of-concept application of Micro-Cm to a single stool specimen, processing larger cohorts will allow optimizing the experimental and data analysis protocol steps and assess reproducibility across individuals and environments.

In summary, Micro-Cm offers an untargeted high-resolution method to explore how a microbial species’ 3D genome changes *in situ* within a complex microbiome community and to perform host-MGE linkage. These advances enable a better understanding of how the interplay between genome topology and function influences community dynamics within complex microbiomes.

## Data availability

The sequencing data associated with the publication have been deposited in the European Nucleotide Archive (ENA); project accession: PRJEB123233. The associated code is available at https://github.com/leylabmpi/microc_paper_code.

## Acknowledgements

This work was funded by Max Planck Society. We thank Stacey Heaver for cultivating bacterial isolates for the simulated defined community.

