## Supplementary figures and methods for "Micro-Cm: restrictase-free microbiome-wide chromosome conformation profiling"

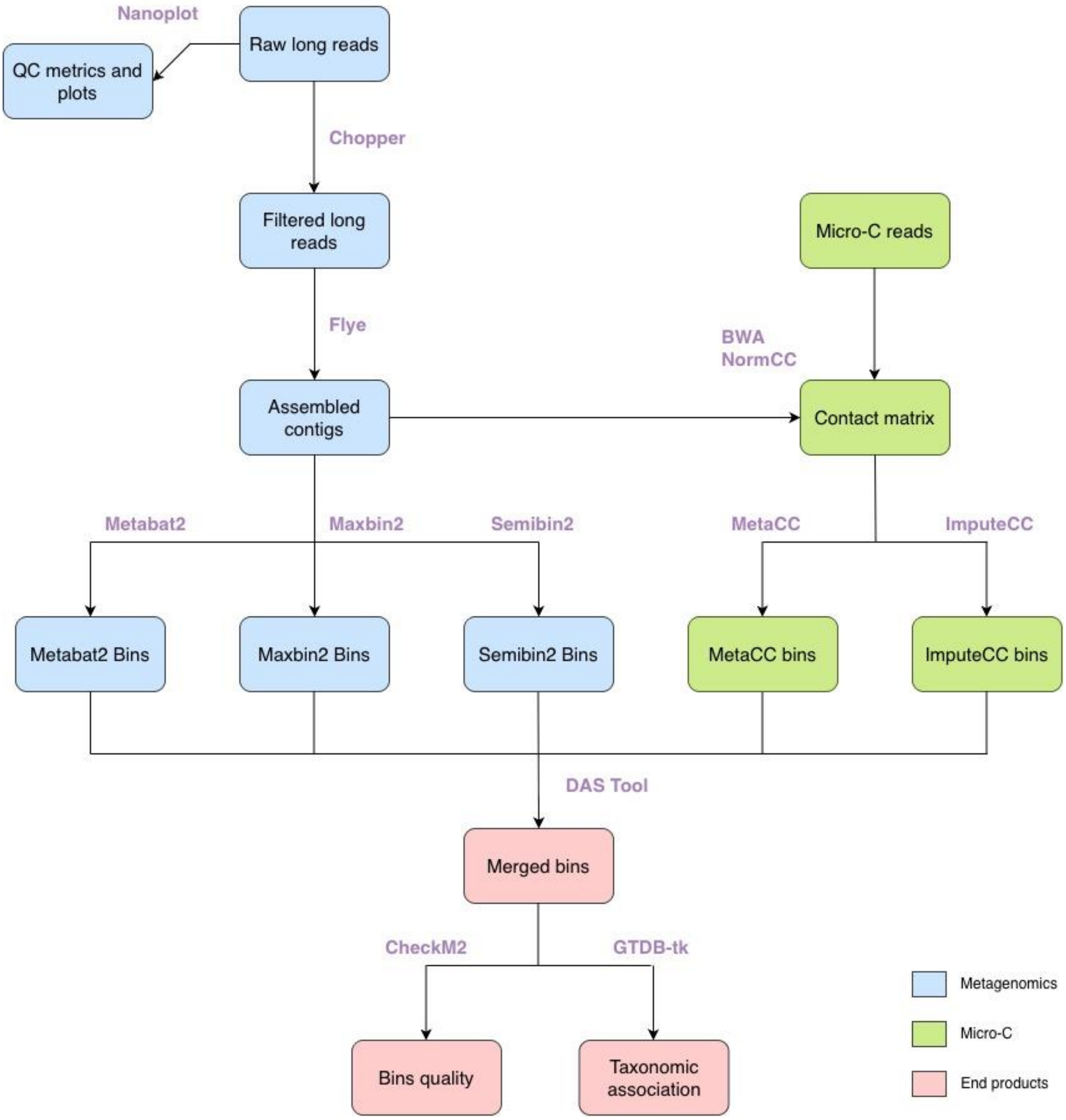


**Suppl. Fig. 1 - Steps involved in the assembly and binning of real microbiome samples.** Long-read metagenomic processes are represented in blue, Micro-Cm-related procedures in green and the final output from the analysis in red. The names of the software employed at each step are specified in purple.

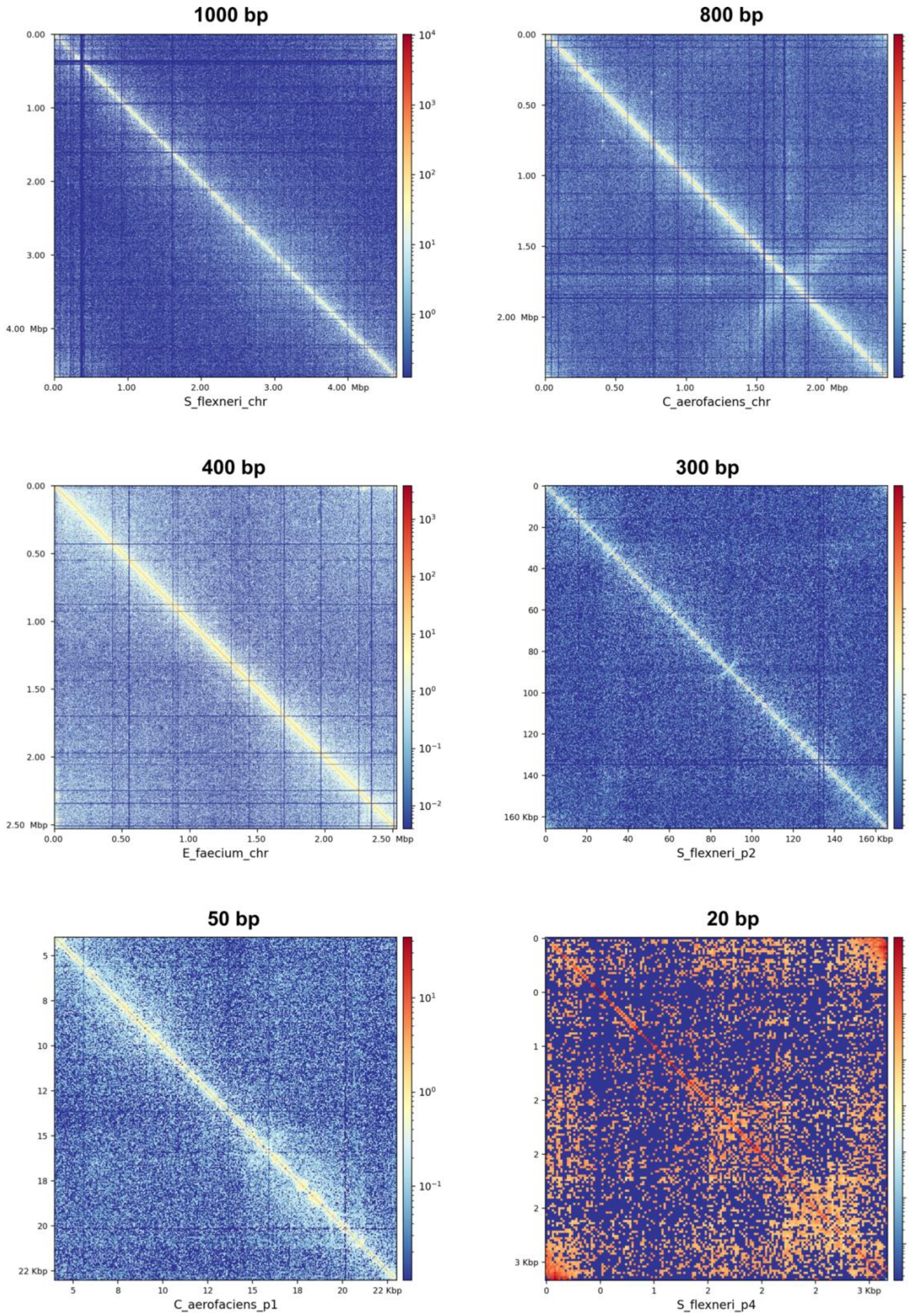


**Suppl. Fig. 2 - Heatmaps showing chromosomal patterns of different sequences in the simulated community.** The resolution of each map is specified above the panel, and the sequence shown is indicated at the bottom. Coordinates are provided on the axes. For *C. aerofaciens* plasmid 1, only part of the sequence is displayed due to partial overlap with the chromosome.

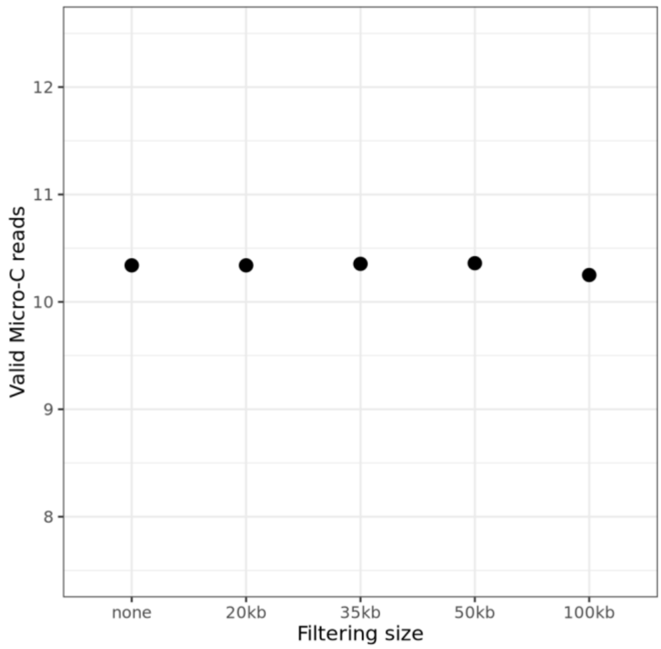


**Suppl. Fig. 3 - Percentage of valid Micro-C reads for the real stool sample after filtering the metagenomic contigs by the various length thresholds specified on the x-axis.**

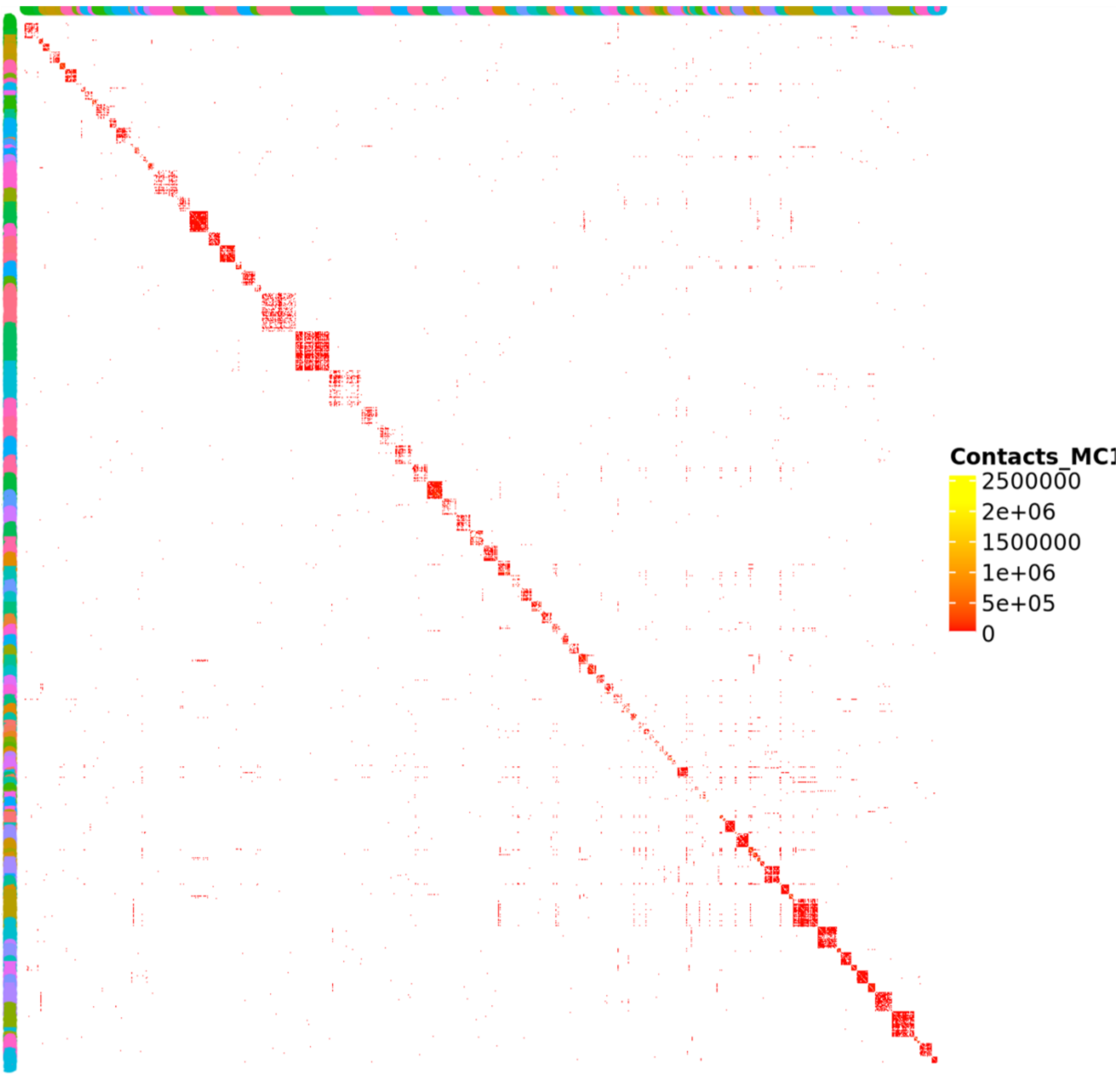


**Suppl. Fig. 4 - Contig-level contact map.** Shown are the high-quality MAGs (n = 101) and only the contacts above the threshold (contact intensities below the threshold were converted to 0). Contigs are grouped by color along the axes according to their bin correspondence.

### Supplementary methods

#### Preparation of modified PVG medium

The DSMZ Medium 104 was prepared by first mixing 5 g/L trypticase peptone, 5 g l/L peptone, 10 g/L yeast extract, 5 g/L beef extract, 5 g/L glucose, 2 g/L $K_2HPO_4$, 1 mL Tween 80, 1 mg resazurin, and 40 mL salt solution. After autoclaving and cooling under an $N_2/CO_2$ atmosphere, 0.2 mL of vitamin $K_1$ solution, 10 mL of hemin solution, and 0.5 g of cysteine-HCl·$H_2O$ were added. The pH was adjusted to 7.2 using 8 N NaOH.

The salt solution was prepared by combining 0.25 g/L $CaCl_2·2H_2O$, 0.5 g/L $MgSO_4·7H_2O$, 1 g/L $K_2HPO_4$, 1 g/L $KH_2PO_4$, 10 g/L $NaHCO_3$, and 2 g/L NaCl. The vitamin $K_1$ solution consisted of 0.1 ml of vitamin $K_1$ dissolved in 20 mL of 95% ethanol, which was then filter-sterilized. Finally, the hemin solution was prepared by dissolving 50 mg of hemin in 1 mL of 1 N NaOH and bringing the final volume to 100 ml with distilled water.

#### Micro-Cm protocol

The first step was to perform formaldehyde crosslinking. For the simulated community, this was done before freezing the sample (see Methods, Simulated community: Bacterial cultivation). For the stool sample, approximately 100 mg of stool was aliquoted and resuspended in a 3 % formaldehyde solution in PBS.

The samples were fixed at room temperature (RT) for 30 min with gentle inversion, quenched with 375 mM Tris-HCl (pH 7.5) and centrifuged at 3,220 × g at 4 °C for 20 min. The pellet was then resuspended in 1 mL of PBS. At this point, the simulated community sample was defrosted and resuspended in 1 mL of PBS. From this point on, both the frozen simulated community and the stool sample were treated in the same way.

All samples were centrifuged at 17,000 × g at 4 °C for 5 min, and the pellet was resuspended in PBS and disuccinimidyl glutarate (DSG, Thermo Fisher) was added to a final concentration of 2 mM (freshly prepared by dissolving in DMSO and subsequently diluting with PBS). The samples were fixed a second time for 45 min at RT with gentle inversion, quenched with 375 mM Tris-HCl (pH 7.5), and centrifuged at 17,000 × g at 4 °C for 5 min.

To disrupt the cells, the pellet was resuspended in 1 mL of 0.9 % NaCl , transferred to prefilled 2.0 mL tubes containing 1.0 mm Zirconium Beads (Biozym), and disrupted on a FastPrep-24 5G (MP Biomedicals) at 6 m/s for 4 × 40 s, with 120 s rest intervals in between. To pellet the cell debris, samples were centrifuged for 10 s at 500 × g, and the supernatant was transferred to a new tube. DNA was pelleted by centrifugation (17,000 × g, 4 °C for 5 min) and resuspended in 1 mL of cold lysis buffer (50 mM Tris-HCl (pH 8), 150 mM NaCl, 0.5 % NP-40 substitute, 1mM EDTA and cOmplete™ Mini EDTA-free Protease Inhibitor Cocktail (1 tablet per 15 mL)) and incubated on ice for 15 min with occasional inversion. Lastly, the samples were centrifuged at 17,000 × g at 4 °C for 5 min, and the pellet was again resuspended in 0.9% NaCl. This washing process was repeated twice.

For the restriction digest, the samples were centrifuged at 17,000 × g for 5 min at 4 °C, and the pellet was resuspended in 300 µL of MNase buffer (10 mM Tris pH 7.5, 1 mM $CaCl_2$). The MNase enzyme (New England Biolabs (NEB), 300 U/µL) was diluted to a final concentration of 2 U/µL using MNase dilution buffer (20 mM Tris pH 7.5, 50 mM NaCl, 50% glycerol) and added to achieve a final amount of 10 U. The reaction mixture was incubated at 37 °C for 25 min with shaking. To inactivate the enzyme, EGTA was added to a final concentration of 5 mM. The samples were centrifuged at 17,000 × g at 4 °C for 5 min, and the pellet was washed in 1× Polynucleotide Kinase (PNK) buffer (NEB) and then resuspended in 145 µL of 1× PNK buffer. 5 µL of PNK (NEB, 10 U/µL) was added, followed by incubation at 37 °C 30 min with shaking. Centrifugation was repeated, and the pellet resuspended in a mix containing 10 µL of 10× NEBuffer r2.1 (NEB), 2 µL of 100 mM ATP, 5 µL of 100 mM DTT and

73 µL water. Finally, 5 µL of PNK were added to each sample, and they were incubated at 37 °C for 15 min with shaking.

The fragmented ends were filled in with biotinylated nucleotides by adding 5 µL of Klenow (NEB, 5 U/µL) and incubation at 37 °C for 15 min with shaking. Subsequently, 5 µL of 10× T4 DNA ligase buffer (NEB), 25 µL of 0.4 mM biotin-dATP (Thermo Fisher), 25 µL of 0.4 mM biotin-dCTP (Thermo Fisher), 1 µL of 10 mM dGTP, 1 µL of 10 mM dTTP and 0.25 µL of BSA (10 mg/ml, NEB) were added, and the mixture was incubated at 25 °C for 45 min with shaking. Afterwards, the samples were centrifuged at 17,000 × g at 4 °C for 5 min, and the pellet was resuspended in 95 µL of 1× T4 DNA ligase buffer. 5 µL of PNK was added, and the solution was incubated at 37 °C and 900 rpm for 1h. Centrifugation was performed as before, and the pellet was resuspended in 478 µL of 1× T4 DNA ligase buffer (Thermo Fisher). 12.5 µL of T4 DNA ligase (5 Weiss U/µl, Thermo Fisher) was added, and the samples were incubated overnight at room temperature while gently rotating.

To remove biotin from unligated ends, the pellet was resuspended in 148.4 µL of 1× NEBuffer 1, and 1.5 µL of exonuclease III (NEB, 100 U/µL) was added. The samples were incubated at 37 °C for 15 min with shaking. The crosslinking was then reversed by adding a mix of 15 µL of 10× NEBuffer 2, 4.5 µL of 5 M NaCl, 30 µL of 10% SDS and 75.5 µL water, after which 25 µL of proteinase K (Thermo Fisher, 20 mg/mL) were added. The mix was incubated at 55 °C for 1h, then for 6h at 65 °C.

DNA was isolated using an equal volume of phenol:chloroform:isoamyl alcohol (25:24:1). It was precipitated using 1/10 volume of 3 M sodium acetate, 3 µL of glycogen and 2.5x volume of 100 % ethanol. Samples were incubated at -80 °C for 40 min, followed by centrifugation at 21,000 × g and 4 °C for 30 min. The DNA was washed with 70% ethanol, air-dried, and then resuspended in 10 mM Tris-HCl (pH 8).

For the RNA digestion, 1 µL of RNase A (Thermo Fisher, 10 mg/mL) was added, and the solution was incubated for 30 min at 37 °C. DNA was purified using 2x AMPure XP beads (Beckman Coulter) and eluted in 40 µL of 10 mM Tris-HCl (pH 8). The volume was then adjusted to 130 µL with sonication buffer (50 mM Tris-HCl, pH 8, 10 mM EDTA, 0.1% SDS).

DNA was sheared using a Covaris S2 (duty cycle 10%, intensity 4, bursts per cycle: 200, 120 sec). Then the samples were transferred to an Amicon 30 K Ultra - 0.5 mL centrifugal filter (Merck) and centrifuged at 16,100 × g and 4 °C for 5 min. This was followed by two washing steps, each consisting of adding 450 µL of 10 mM Tris (pH 8) and centrifuging with the same settings. The concentrated samples were transferred to a new LoBind tube and brought to 100 µL with 10 mM Tris (pH 8).

Biotin pulldown was performed using Dynabeads Streptavidin C1 beads (Thermo Fisher). They were washed with 400 µL of Tween washing buffer (TWB; 5 mM Tris pH 8, 0.5 mM EDTA, 1 M NaCl, 0.05% Tween 20). The beads were resuspended in 100 µL of binding buffer (10 mM Tris pH 8, 1 mM EDTA, 2 M NaCl), the 100 µL of DNA were added and incubated at RT for 20 min while rotating. The beads were then washed: twice with 600 µL of TWB, once with 100 µL 1× NEBuffer 2, and once with 100 µL 1× T4 DNA ligase buffer (NEB). The DNA ends were repaired by resuspending in 75 µL of end repair solution: 7.5 µL of T4 DNA ligase buffer, 1.9 µL of 10 mM dNTP mix, 5 µL of PNK, 1 µL of Klenow, 4 µL of T4 DNA polymerase, and 55.6 µL of nuclease-free water and incubated for 30 min at 22 °C.

For the A-tailing, beads were washed twice with 600 µL of TWB, once with 100 µL 1× NEBuffer 2, and then resuspended in a mixture of 1× NEBuffer 2 and 0.21 mM dATP. 5 µL of Klenow (exo-) (NEB, 5 U/µL) was added, and the reaction mixture was incubated at 37 °C for 40 min. To proceed with the adapter ligation, the beads were washed: twice with 600 µL of TWB, once with 1× NEBuffer 2, and once with 1× T4 DNA ligase buffer (NEB). They were then resuspended in 46.5 1× T4 DNA ligase buffer (NEB), and 2.5 µL of IDT for Illumina TruSeq RNA UD indexes v2 were added. Adapters were ligated for 2.5 h at 22 °C, with occasional mixing, followed by three washing steps: once with 600 µL TWB, once with 100 µL 1× NEBuffer 2, and once with 100 µL of 10 mM Tris HCl (pH 8). Then, the final product was resuspended in 16 µL of nuclease-free water.

4 µL of streptavidin-bound DNA was mixed with 1x KAPA HiFi Buffer, 0.3 mM dNTP mix, 0.5 µM Illumina forward primer, 0.5 µM Illumina reverse primer, and 1 U of KAPA HiFi

DNA Polymerase (4 reactions per sample). PCR amplification was performed with the following program: 95 °C for 5 min, then 13 cycles of 98 °C for 20 sec, 65 °C for 15 sec, and 72 °C for 20 sec, and finally 72 °C for 3 min.

After the PCR, the 4 reactions per sample were pooled, cleaned up using 0.85x AMPure XP beads, and paired-end sequenced on a Illumina NextSeq 2000 platform.
